# Heterogeneity of Inflammation-associated Synovial Fibroblasts in Rheumatoid Arthritis and Its Drivers

**DOI:** 10.1101/2022.02.28.482131

**Authors:** Melanie H Smith, Vianne R Gao, Michail Schizas, Alejandro Kochen, Edward F DiCarlo, Susan M Goodman, Thomas M Norman, Laura T Donlin, Christina S Leslie, Alexander Y Rudensky

**Affiliations:** Division of Rheumatology, Department of Medicine, Hospital for Special Surgery, New York, NY 10021, USA; Howard Hughes Medical Institute and Immunology Program at Sloan Kettering Institute, Ludwig Center for Cancer Immunotherapy, Memorial Sloan Kettering Cancer Center, New York, NY 10065, USA; Computational and Systems Biology Program, Memorial Sloan Kettering Cancer Center, New York, NY 10065, USA; Arthritis and Tissue Degeneration Program and the David Z. Rosensweig Genomics Research Center, Hospital for Special Surgery, New York, NY 10021, USA; Department of Pathology and Laboratory Medicine, Hospital for Special Surgery, New York, NY 10021, USA; Weill Cornell Medical College and Graduate School, New York, NY 10021, USA

## Abstract

Inflammation of non-barrier immunologically quiescent tissues is associated with a massive influx of blood-borne innate and adaptive immune cells. Cues from the latter are likely to alter and expand the spectrum of states observed in cells that are constitutively resident. However, local communications between immigrant and resident cell types in human inflammatory disease remain poorly understood. Here, we explored heterogeneity of synovial fibroblasts (FLS) in inflamed joints of rheumatoid arthritis (RA) patients using paired single cell RNA and ATAC sequencing (scRNA/ATAC-seq), multiplexed imaging, and spatial transcriptomics along with *in vitro* modeling of cell extrinsic factor signaling. These analyses suggest that local exposures to myeloid and T cell derived cytokines, TNFα, IFNγ, IL-1β, or lack thereof, drive six distinct FLS states some of which closely resemble fibroblast states in other disease-affected tissues including skin and colon. Our results highlight a role for concurrent, spatially distributed cytokine signaling within the inflamed synovium.

---

Rheumatoid arthritis (RA), a systemic autoimmune disease with predominantly articular manifestations, is characterized by hyperplasia of both the synovial lining, which interfaces the synovial sublining and the synovial fluid-filled joint space, as well as the synovial sublining, which exhibits increased vascularization and an influx of leukocytes. Both the lining and sublining fibroblast-like synoviocytes (FLS) undergo proliferation and activation, assuming a state, in which they stimulate angiogenesis, produce pro-inflammatory cytokines and chemokines, and invade adjacent articular cartilage and bone^1^. Expression of MHC class II molecules by activated FLS is associated with synovial inflammation^2,3^ and their expression by lining FLS correlates with active disease^4^. HLA-DR^+^ FLS expression of soluble mediators, including the proinflammatory cytokines IL-6 and IL-15, and chemokines CCL2, CXCL9, and CXCL12 along with adhesion molecules such as ICAM1 and VCAM1 suggests that these features of FLS might be imparted by their interactions with leukocytes^2^. In support of this possibility, prior *in vitro* studies have shown that HLA-DR^+^ FLS are capable of presenting antigens to CD4^+^ T cells^5–7^. Furthermore, production of the aforementioned proinflammatory chemokines by FLS likely acts as a feedforward mechanism to further facilitate recruitment of diverse immune cell types expressing the corresponding receptors. Indeed, a recent study of the overall cellular makeup of synovial tissue from RA patients using single cell RNA sequencing (scRNA-seq) analysis identified a diverse mix of migratory and resident cell types of hematopoietic and non-hematopoietic origin including different CD4 and CD8 T cell subsets, myeloid cells, and FLS^2^. These observations suggest that states of FLS activation and differentiation are likely modulated by diverse innate and adaptive immune cell types found in inflamed joints of RA patients and that this modulation factors prominently in the disease pathogenesis.

Thus, we sought to undertake an in-depth investigation of the spectrum of FLS states induced in the inflamed RA synovium and potential drivers underlying the observed FLS heterogeneity through paired scRNA and assay for transposase-accessible chromatin with sequencing (scRNA/ATAC-seq) and *in vitro* modeling of FLS transcriptional responses to key immune cell-derived proinflammatory cytokines. We then mapped the spatial distribution of FLS heterogeneity and transcriptional responses by employing spatial transcriptomic (ST) analyses as well as multiplex imaging. Our studies suggest that spatially constrained FLS responses to three major leukocyte-derived cytokines, TNFα, IFNγ, and IL-1β, or lack thereof, drive the formation of six distinct FLS states associated with synovial inflammation in RA.

## Results

### Inflammation is associated with heightened FLS heterogeneity

To test the possibility that severe joint inflammation in RA patients leads to a marked expansion of FLS heterogeneity, we sought to characterize their transcriptomes and chromatin accessibility at a single cell resolution. Fluorescence-activated cell sorting (FACS)-sorted FLS (CD45^-^CD31^-^PDPN^+^) were isolated from two RA patients with highly inflamed synovium, who had similar disease characteristics as well as histologic findings, and subjected to paired scRNA/ATAC-seq using the 10x Multiome platform (patients 1 and 2 in Table S1, Extended Data Fig. 1a). After extensive filtering, we obtained 15,736 FLS that clustered into 14 clusters based on the scRNA-seq datasets (Fig. 1a, Extended Data Fig. 1b-e, Table S2). Similar results were obtained with and without Mutual Nearest Neighbors (MNN) batch correction (Extended Data Fig. 1f,g). Using established characteristic markers^2^, some of which are shown in Fig. 1b, we identified lining, sublining and pan-synovial clusters. The latter clusters express genes characteristic of both sublining and lining FLS. Consistent with the high degree of synovial inflammation, MHC class II expression was widespread as HLA-DR expressing cells were found in all clusters except for cluster 13 (Fig. 1c). Our observation of a high fraction of lining FLS expressing HLA-DR was at odds with previous reports that HLA-DR^+^ FLS represent a subset within the sublining FLS population^2^. Therefore, we independently assessed HLA-DR expression using multicolor immunofluorescence (IF) and confirmed the pan-synovial expression of HLA-DR on FLS (Fig. 1d).

**Figure 1.**
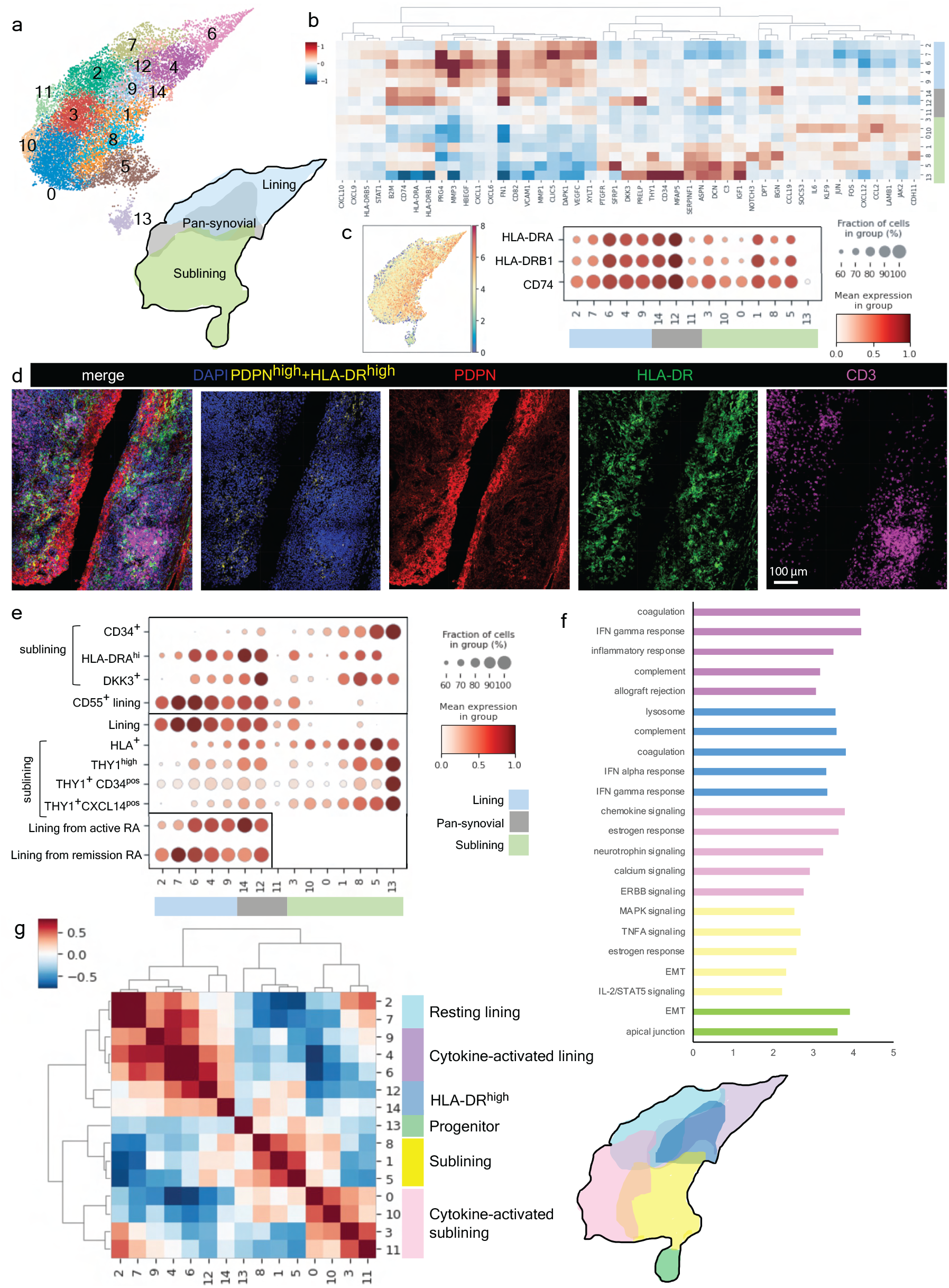
Heterogeneity of HLA-DR^+^ FLS in the inflamed RA synovium. **a,** UMAP of 14 FLS clusters identified by scRNA-seq analysis with annotations of synovial localization. **b,** Heatmap of selected DEGs for each cluster colored by synovial localization. **c,** UMAP colored by logged, library size-normalized expression of HLA-DRA and dot plot showing relative expression of HLA-DRA, HLA-DRB1 and CD74. **d,** Representative confocal microscopy of PDPN (red), HLA-DR (green), CD3 (magenta) and nuclear marker (blue) from RA synovial tissue (N = 5 tissues). Pixels with the highest intensity (top 3%) for both PDPN and HLA-DR are colored in yellow. **e,** Dotplot showing the relative per cluster expression of previously published cluster-derived gene signatures: Zhang et al^2^ above and Alivernini et al^4^ below horizontal line. Box (bottom left) shows lining FLS signatures from patients in remission or with active disease from Alivernini et al. **f,** GSEA showing top 5 pathways from KEGG with FDR <0.1 for each of the states defined in G. The resting lining state did not have any statistically enriched pathways. **g,** Cluster by cluster correlation of the mean expression of highly variable genes in clusters from Fig. 1a with defined FLS states colored on UMAP.

In addition to capturing all FLS subsets previously identified in the RA synovium^2,4^ (Fig. 1e), our focused analysis of FLS in highly inflamed tissue enabled identification of novel subsets of both sublining (clusters 0, 10, 11) as well as lining (clusters 4, 6, 9) FLS (Fig. 1e). The large number of lining FLS spanning multiple clusters included those sharing transcriptional signatures characteristic of FLS derived from patients with both active and remission RA^4^. These results suggest that a highly inflamed RA joint harbors a greatly expanded range of FLS states possibly representing a broad spectrum of the disease.

While each cluster had specific features, such as the expression of *NOTCH3* in cluster 8 that marks perivascular FLS^8^, gene set enrichment analysis (GSEA) of each cluster highlighted shared functionality between groups of clusters (Fig. 1f and Table S3). In fact, the assessment of cluster-by-cluster correlation, allowed us to define six FLS states each with distinct inferred functionality (Fig. 1g). The identified resting lining FLS state shared a transcriptional profile with lining FLS from RA patients in remission^4^. The cytokine-activated lining FLS state, which expressed elevated levels of HLA-DR, displayed an IFNγ response gene signature and additional inflammatory response gene signatures. Pan-synovial HLA-DR^high^ FLS were characterized by the highest expression of HLA-DR, and also displayed an IFNγ response signature as well as lysosome/endosome-related genes (*CD63, CTSD, NPC2, LAPTM4A, CTSK, CD68, CTSL, CTSA, HEXA, CTSF, GUSB, LAMP1*) suggesting their potential role as antigen-presenting cells in the inflamed synovium. A transcriptional profile of these pan-synovial HLA-DR^high^ FLS closely resembled the one previously reported for lining FLS from patients with active RA^4^. The sublining FLS were enriched for cytokine signaling pathway genes, most notably TNFα, but also IFNγ and IL-6. Of note, both the sublining and cytokine-activated sublining FLS states show evidence of STAT5 signaling possibly driven by IL-15. Finally, we observed that the transcriptomes of a subset of sublining FLS exhibited features expected for progenitor cells including the highest level of CD34 expression, and enrichment for expressed genes related to extracellular matrix (ECM) homeostasis and epithelial mesenchymal transition (EMT) (*MFAP5, FBN1, VCAN, TGFBR3, FBLN2, PRRX1, DCN, LAMA2, SFRP4, EDIL3, FBLN1, LGALS1, LOXL1, ADAM12, LOX, CD44, IGFBP4, LRP1).* Notably, the most differentially expressed gene for the progenitor state was PI16, which was recently described as a marker for one of two populations of universal mouse (“cross-tissue”) fibroblasts that can give rise to specialized fibroblast populations during development and upon perturbation^9^. Furthermore, the gene expression features of the progenitor FLS state we characterized showed extensive similarity to the PI16^+^ cluster and enrichment for the “universal fibroblast gene signature” identified in a human perturbed-state fibroblast atlas (Extended Data Fig. 1h). Since tissue progenitor or “stem-like” cells can be frequently found as aggregates within distinct specialized anatomical niches commonly associated with vasculature, we explored spatial distribution of these CD34^high^THY1^+^PDPN^+^ FLS using IF. However, we found them widely dispersed as solitary cells throughout the inflamed synovium without conspicuous association with the vascular endothelium (Extended Data Fig. 1i). This finding suggests a possibility that in highly inflamed RA synovium FLS regenerative capacity is preserved in a non-compartmentalized manner.

### Comparison with non-synovial fibroblasts shows shared functionality across tissues

Previous cell population-based studies suggested distinct diversity of transcriptional features of human fibroblasts in different anatomical locations and heritable imprinting of their “topography”^10^. However, our observation of conserved “universal fibroblast” features of progenitor PI16^+^ FLS suggested that there might be an overlap between disease-induced states of anatomically distinct tissue fibroblasts affected by different pathologies. Thus, we next sought to explore whether other FLS states we identified were tissue or disease-specific, namely, unique to the synovium or RA-associated inflammation, or alternatively, were shared with fibroblast populations observed in other diseases and tissues (Fig. 2, Extended Data Fig. 2). For this comparative analysis, we took advantage of several recent scRNA-seq datasets of colonic^11,12^ and dermal fibroblasts^13,14^, in which at least 5 fibroblast clusters can be delineated.

**Figure 2.**
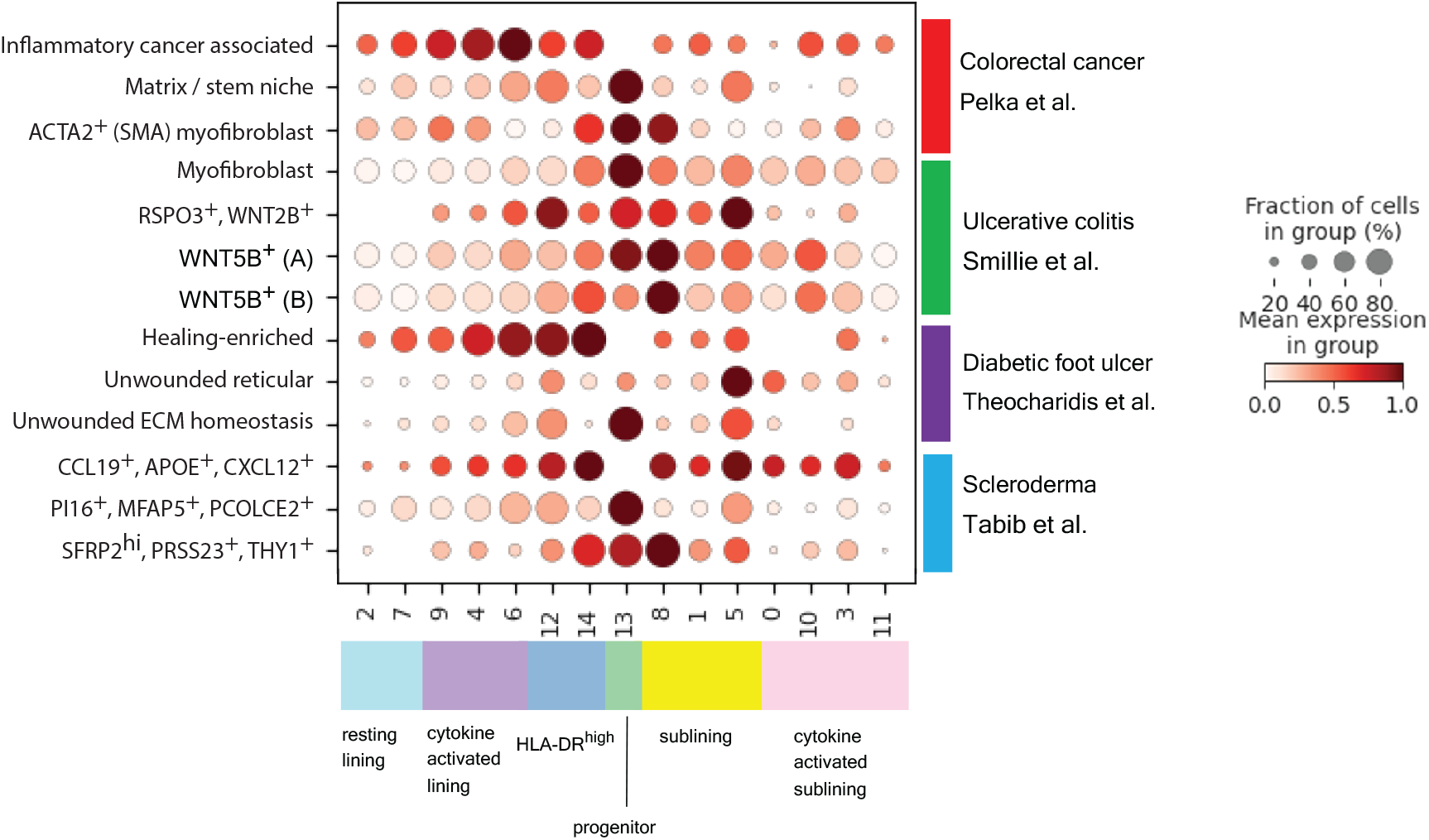
Shared functional gene expression programs in FLS and non-synovial fibroblasts across tissues and diseases. Dot plot showing relative expression of selected gene signatures from published tissue fibroblast populations^11–14^ in FLS clusters shown in Fig. 1a colored according to FLS states defined in Fig. 1g.

As expected from the aforementioned presence of the “universal fibroblast signature”, the progenitor FLS state shared transcriptional similarity with fibroblasts from both the colon and skin. In the colon, this transcriptional signature was observed in two fibroblast populations implicated in creating an intestinal stem cell niche: ECM producing fibroblasts and myofibroblasts. In the skin, this transcriptional signature was observed in fibroblasts from unwounded skin responsible for ECM homeostasis and in a fibroblast population that undergoes contraction in scleroderma as compared to healthy skin.

Unexpectedly, we observed a remarkable transcriptional similarity between inflammatory cancer associated fibroblasts in colorectal cancer and the HLA-DR^+^ cytokine-activated lining FLS. The latter FLS state was also similar to transcriptional states of fibroblasts in diabetic foot ulcers that were able to successfully heal.

On the other hand, transcriptional features of fibroblasts in the diseased skin or colon bore only limited similarity to both the resting lining and cytokine-activated sublining FLS states we observed. The resting lining FLS were characterized by high expression of genes involved in the production of synovial fluid and ECM (*XYLT1, FN1, ITGB8, PRG4*) and axonal guidance (*SEMA5A, ANK3, NTN4, SLIT2),* features which likely reflect their specialized functions within the synovium. The cytokine-activated sublining FLS state also appeared distinct to the RA synovium, however, it likely emerged due to the high degree of inflammation in the RA synovial samples we analyzed. Accordingly, corresponding fibroblast states in other tissues are expected to be found only under similarly highly inflamed conditions and may not have been enriched in the scRNA-seq datasets used for the cross-comparison. Thus, inflammation-induced perturbations in the overall composition of the FLS population and spectrum of FLS states were shared with fibroblasts found in other tissues in a range of pathologies.

### FLS states exhibit distinct transcriptional regulation with evidence of differential cytokine stimulation

To gain insight into transcription factors and upstream signaling pathways which promoted FLS heterogeneity in RA, we analyzed paired scATAC-seq datasets. Unsupervised clustering resulted in seven clusters with lining and sublining FLS subtypes as well as an independent cluster of progenitor-like FLS (Fig. 3a,b, Extended Data Fig. 3a-c). Distinct FLS states identified by scRNA-seq analyses occupied divergent areas of the scATAC-seq UMAP (Fig. 3c) partially overlapping with the identified scATAC-seq clusters (Extended Data Fig. 3d). To infer differential transcription factor activity in identified FLS states, we performed chromatin accessibility variation analysis using chromVAR. For this analysis, we used the paired scRNA-seq data as a filter to assess only motifs for the transcription factor families whose members were expressed by >20% of cells in the corresponding state. We observed marked differences in enrichment of distinct transcription factor binding motifs within open chromatin sites with differential motif accessibility between states (Fig. 3d,e). The cytokine-activated lining and HLA-DR^high^ pan-synovial FLS states were enriched for activity of AP-1 transcription factor family members (JUN, JUNB, JUND, FOS, FOSL2), whose increased contribution to gene regulation downstream of fibroblast growth factor (FGF) and immune receptor signaling, such as IL-1, has been suggested to play a role in tissue-destructive properties of FLS in RA^15,16^. Interestingly, open chromatin sites characteristic of the cytokine-activated sublining FLS state were enriched for STAT and IRF family motifs implicating a distinct set of inflammatory pathways such as IFN signaling in establishing this state. Contrary to the two major flavors of activated, inflammation-associated FLS, the resting lining FLS state was distinguished by the accessibility of homeobox transcription factor family member motifs, which besides serving as major regulators of development and organization, control fibroblast quiescence (e.g., PRRX1)^17,18^. In support of a role for homeobox transcription factors in modulating FLS activation, the resting lining FLS state exhibited increased expression of homeobox family member *CUX1,* which has been shown to bind to NF-kB and alter its activity through either downregulation^19^ or upregulation^20^ of specific NF-kB-regulated cytokines and chemokines. Finally, the PI16^+^ progenitor FLS were distinguished by accessible *cis*-regulatory elements enriched for KLF, SOX and TEAD family motifs, which play a role in maintaining quiescent undifferentiated states in stem cells and early progenitors. In this regard, KLF4 has been implicated in the induction of a pluripotent state in fibroblasts consistent with the likely role of this FLS subset as progenitors^21^. These results suggest that distinct states of FLS in the inflamed RA joint were dependent upon their local stimulation by immune cell derived factors, foremost proinflammatory cytokines, or avoidance of these inflammatory exposures.

**Figure 3.**
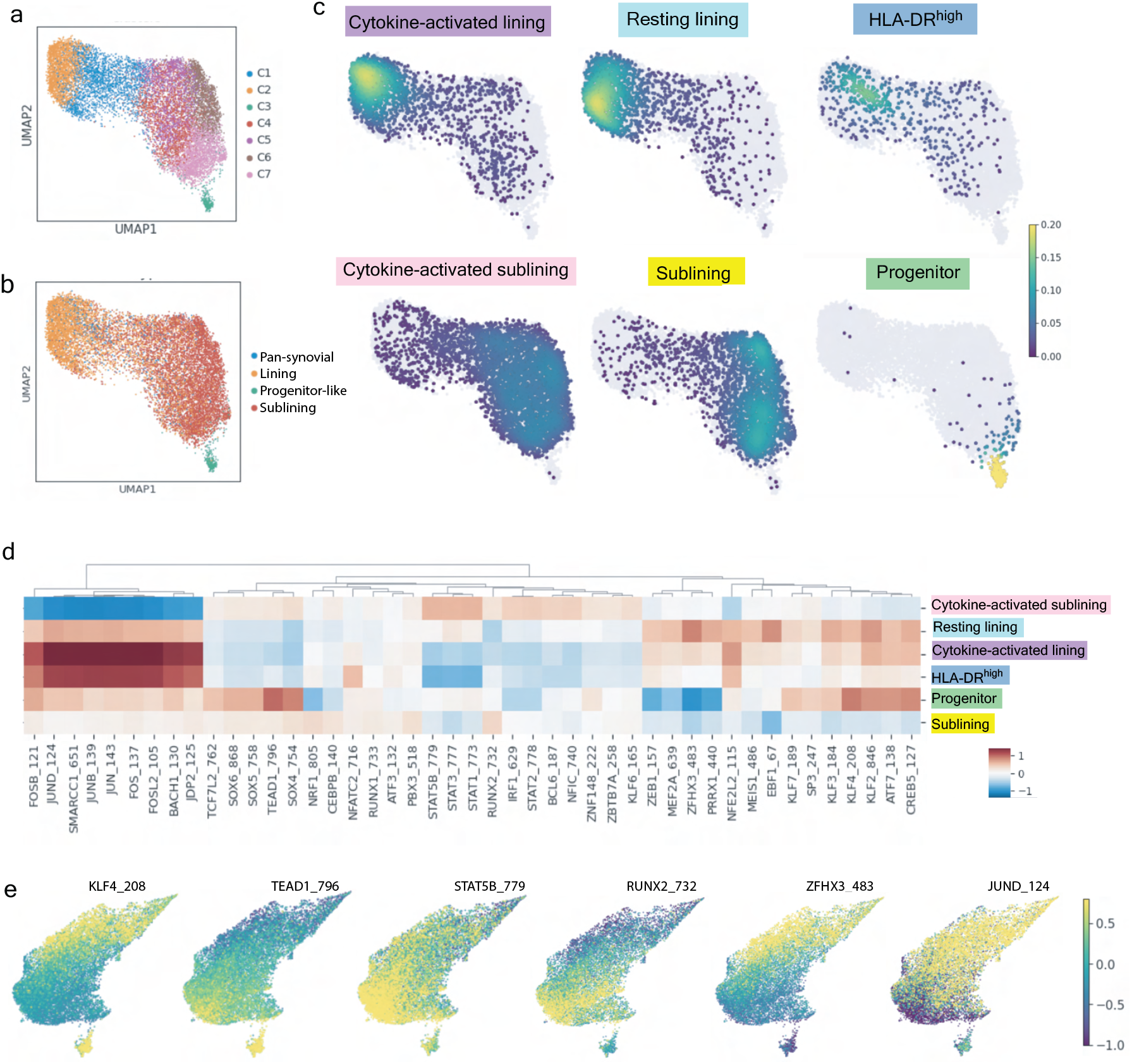
Chromatin accessibility analysis of FLS states reveals their distinct transcriptional regulation. **a,** UMAP of 7 FLS clusters identified by tile-based scATAC-seq analysis. **b,** Annotations of synovial localization as well as the progenitor state on the scATAC-seq UMAP. **c,** Projection of six FLS states onto scATAC-seq UMAP. **d,** Heat map with top 10 differentially accessible transcription factor motifs identified by ChromVAR for each FLS state defined in Fig. 1g. Motifs filtered to include only those for which the corresponding transcription factor was expressed by >20% of cells in the corresponding state. **e,** ChromVAR z-score projected onto scRNA-seq UMAP for a selection of top differentially accessible transcription factor motifs derived from each FLS state.

### Cytokine signaling drives transcriptional heterogeneity

To test the above possibility and to elucidate influences of inflammatory factors on transcriptomes of FLS states we sought to deconvolute their complex transcriptomes by establishing cell-type specific cytokine-induced programs. For identification of immune cell derived cytokine transcriptional responses and their contributions to distinct features of FLS state-specific transcriptomes, we employed *in vitro* stimulation of cultured FLS by candidate pro-inflammatory cytokines and other factors. For these experiments, FLS were isolated from four RA synovial tissue samples and cultured for three passages prior to pooling FLS from all donors and performing the stimulations in triplicate. FLS were stimulated with three major cytokines implicated in RA – TNFα, IFNγ and IL-1β, either individually or in combination – and the resulting gene expression changes were assessed using RNA-seq (Fig. 4a, Table S4). Additionally, to directly compare cytokine stimulation effects on a per patient basis, we isolated FLS from the same RA synovial tissue samples subjected to scATAC/RNA-seq (Figs. 1 and 3) stimulated them with TNFα, IFNγ and IL-1β or TNFα and IFNγ and performed scATAC/RNA-seq. We found that *in vitro* stimulation of FLS with both TNFα and IFNγ induced expression of genes including *CCL2* and *IL6,* which were highly expressed by *ex vivo* isolated cytokine-activated sublining FLS. On the other hand, genes whose expression was markedly suppressed in response to these cytokines (e.g., *VCAN, CCDC80, CD248* in Fig. 4a) were highly upregulated by the CD34^high^THY1^+^PI16^+^ FLS suggesting that the progenitor FLS state is shielded from exposure to inflammatory mediators and that these cytokines may lead to the loss of this state. In the same vein, *ANK3,* which is upregulated in the resting lining FLS state, was also downregulated in response to *in vitro* stimulation by TNFα and IFNγ suggesting that besides the progenitor FLS in the sublining, resting lining FLS also appear to be spared from the full effects of inflammatory cytokines. Finally, genes induced in FLS subjected to *in vitro* stimulation by the combination of TNFα, IFNγ and IL-1β, which included *MMP3* and *CXCL1,* were most differentially expressed in *ex vivo* isolated cytokine-activated lining FLS.

**Figure 4.**
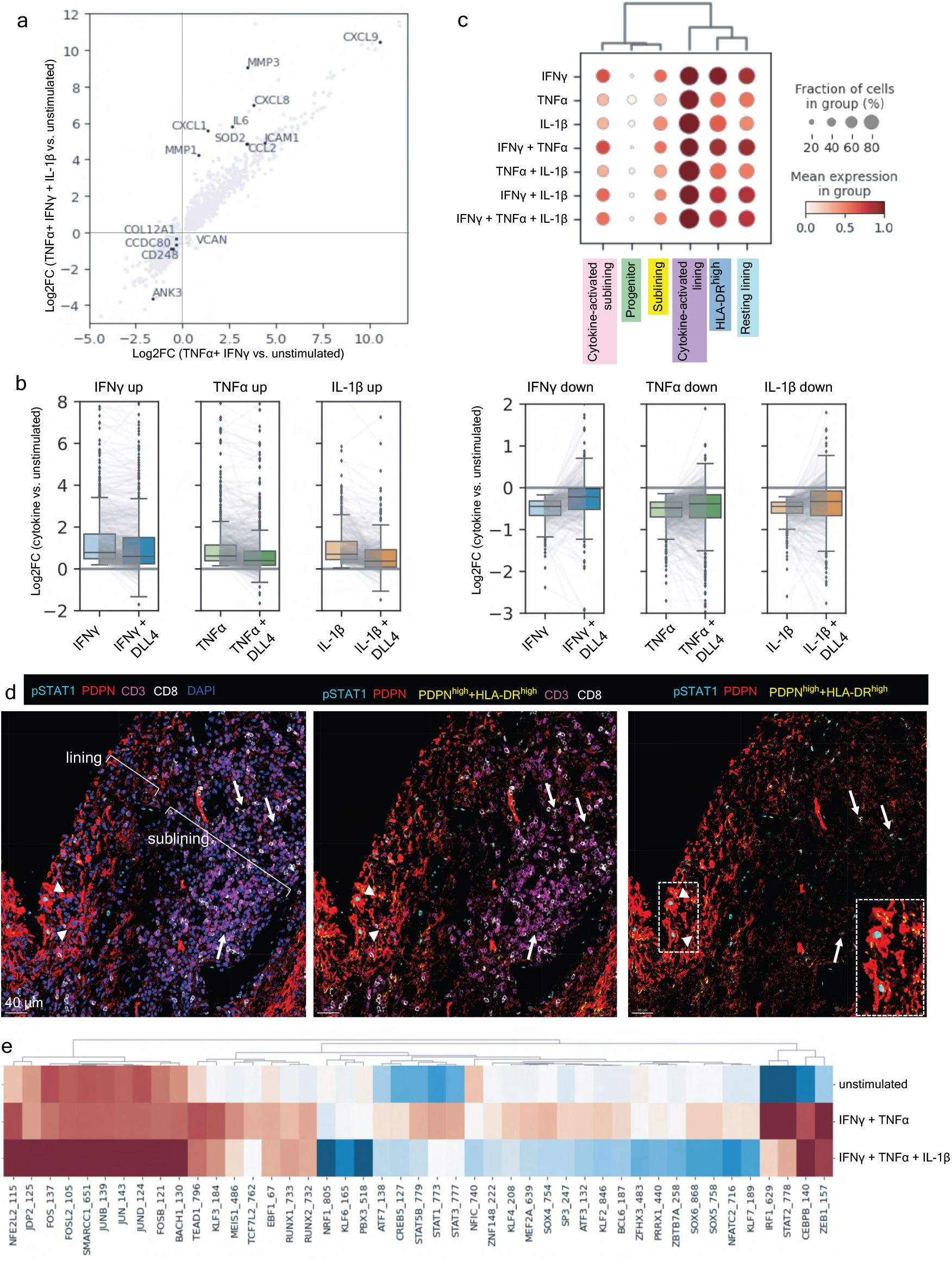
Cytokine signaling drives transcriptional FLS heterogeneity. **a,** Changes in gene expression (log fold change) after combinatorial stimulation of cultured FLS by cytokines indicated. **b,** Cultured FLS were treated with the individual cytokines indicated *in vitro* and subjected to 3’ RNA sequencing to identify genes that were up- (left) or down- (right) regulated. Box plots compare the distribution of log2 fold changes in the expression of these genes (in stimulated versus control) in FLS treated with each cytokine alone or in combination with DLL4. Gray lines connect individual genes across conditions. **c,** Dot plot showing relative expression of the identified cytokine response signatures in each of the FLS states defined in Fig. 1g. **d,** Representative confocal images of phosphorylated STAT1 staining (cyan), PDPN (red), CD3 (magenta), CD8 (white) and nuclear marker (blue) (N = 4 tissues). Pixels with the highest intensity (top 3%) for both PDPN and HLA-DR are colored in yellow. White arrows indicate FLS with nuclear pSTAT1 staining in the sublining and white arrowheads indicate FLS with nuclear pSTAT1 staining in the lining. **e,** ChromVAR z-score of motifs from Fig. 3d in cultured FLS that were unstimulated, simulated with TNFα + IFNγ, or stimulated with TNFα, IFNγ and IL-1β.

Notch signaling induced by ligands expressed by the vascular endothelium have been suggested to factor prominently in the differentiation of perivascular and sublining FLS in the RA synovium^8^. This raised the question as to whether Notch signaling can potentially modulate transcriptional responses of FLS to pro-inflammatory cytokines within the sublining. To explore this possibility, we investigated changes in gene expression induced in FLS upon stimulation with TNFα, IFNγ and IL-1β in the presence or absence of plate-bound Notch ligand Delta like-4 (DLL4). This analysis revealed a global dampening of transcriptional responses to all three cytokines (Fig. 4b, Table S5): for both up- and down-regulated genes, the observed changes were blunted across the board. DLL4 similarly attenuated transcriptional responses to dual and triple combinations of cytokines (TNFα + IFNγ and TNFα + IFNγ + IL-1β, respectively) (data not shown). This finding was unexpected considering that previous studies suggested that Notch signaling augments macrophage responses to TLR ligands and increases production of pro-inflammatory cytokines^22^. However, there may be multiple specific regulatory mechanisms in play as previous reports also show that IFNγ can inhibit Notch signaling in macrophages^23^. These results raise an intriguing question as to whether coincident engagement of inflammatory and developmental signaling pathways may result in different functional outcomes depending on a given cell type.

Mapping of cytokine response gene signatures established by the above *in vitro* analyses onto our scRNA-seq datasets revealed the most pronounced expression of these gene signatures in the cytokine activated lining FLS state (Fig. 4c). In contrast, the dominant cytokine response signatures in the cytokine-activated sublining FLS state included either genes modulated by a dual combination of TNFα and IFNγ or by IFNγ alone. Notably, the progenitor state was devoid of cytokine response gene signatures.

Previous scRNA-seq analysis suggests that CD8^+^ T cells represent the major source of IFNγ in the RA synovium^2^. In agreement with previous reports, our imaging showed predominant localization of CD8^+^ T cells within lymphoid aggregates in the sublining region. Therefore, we sought to investigate whether local FLS interferon responses correlated with CD8^+^ T cell localization *in situ* by analyzing phosphorylated STAT1 (pSTAT1), the key downstream target of interferon signaling, using IF (Fig. 4d). Interestingly, we observed nuclear pSTAT 1 in PDPN^+^ FLS within both lymphocyte aggregates in the sublining region as well as the T cell poor lining region. Of note, some of the PDPN^+^pSTAT1^+^ FLS also express HLA-DR consistent with a well-recognized role of IFNγ in driving MHC class II expression, while pSTAT1 observed in T cell poor regions may reflect local type I IFN signaling.

To further validate the effect of cytokine stimulation on regulation of gene expression, we performed scATAC-seq of cultured FLS stimulated with combinations of cytokines to cross-reference the observed transcription factor motif activity at modulated chromatin accessibility sites to that of distinct FLS states revealed by scATAC-seq analyses of *ex vivo* isolated cells (Fig. 4e). We found that stimulation of FLS with a combination of TNFα and IFNγ resulted in enrichment of IRF and STAT family transcription factor motifs at differentially accessible chromatin sites in comparison to unstimulated FLS, closely matching those in the cytokine-activated sublining state of FLS *ex vivo.* In contrast, triple combination of TNFα, IFNγ and IL-1β resulted in differential chromatin remodeling at sites enriched for AP-1 transcription factor family motifs. The latter observation was consistent with a previous report of remodeling of chromatin regions containing NF-kB and AP-1 binding motifs in response to IL-1β^24^. Impressively, the differentially accessible sites enriched for AP-1 family motifs were nearly identical to those observed in the *ex vivo* isolated cytokine-activated lining FLS state. The comparison of cytokine-stimulated samples to the unstimulated control explains the seemingly unexpected decrease in accessibility of *cis*-regulatory elements containing STAT motifs in the cytokine-activated lining FLS (Fig. 3d) despite the presence of a robust IFNγ response gene expression signature (Fig. 4c) and STAT1 phosphorylation (Fig. 3d) (Extended Data Fig. 4). Thus, this was most likely due to a relative decrease in STAT accessibility of a subset of these elements caused by combined IL-1β, IFNγ and TNFα exposure while the corresponding transcript levels were not markedly impacted. Together, these analyses of transcriptomes and epigenomes strongly support a role of combinatorial stimulation by TNFα and IFNγ in facilitating the establishment of the cytokine-activated sublining FLS state whereas the cytokine-activated lining FLS state was likely driven by a triple combination of TNFα, IFNγ and IL-1β.

### Cytokine signaling is spatially constrained and correlated with cellular localization

The observations above suggest that distinct states of FLS in the inflamed synovium are established in a spatial manner as the result of locally produced inflammatory cytokines and other mediators by distinct types of immune cells invading the RA synovium. To define the spatial distribution of identified FLS states within the RA synovium, we performed spatial transcriptomic (ST) analyses using the 10x Visium platform combined with multiplex IF imaging of adjacent tissue sections for two additional inflamed synovial tissue samples isolated from RA patients (patients 3 and 4 in Table S1). The hematoxylin and eosin (H&E) staining of the sections subjected to ST analysis showed prominent lymphocyte aggregates as well as copious synovial lining (Fig. 5a). IF imaging of the sections adjacent to those used for ST showed scattered PDPN^+^ FLS and CD68^+^ macrophages as well as lymphocyte aggregates in the sublining region, and multiple regions of lining populated by FLS and macrophages (Fig. 5b). The ST datasets were integrated with our scRNA-seq analyses to map the transcriptional signatures from the six FLS states on the ST datasets. This showed that the resting and cytokine-activated lining FLS appeared to intermix without forming well-defined regions. In contrast, the three identified sublining FLS subsets were differentially localized (Fig. 5c). We next applied the *in vitro* FLS cytokine response gene signatures to spatial gene expression maps derived from ST analysis. We found that the IL-1β response signature mapped predominantly to the synovial lining, while the other cytokine response signatures were more scattered (Fig. 5d). Since multiple cells contribute to each RNA capture spot on a Visium slide, we confirmed that the IL-1β response gene signature observed in ST analysis was contributed by FLS rather than other cell types by creating a modified IL-1β response gene signature that only contained genes uniquely expressed by FLS based on recent scRNA-seq analysis of RA synoium^2^. Integrated ST and IF imaging analysis of adjacent sections showed that the cytokine-activated lining FLS state featuring the IL-1β response gene signature was near areas densely populated by CD68^+^ macrophages. Mapping cell type specific transcriptional signatures from a recent scRNA-seq analysis of the RA synovium^2^ onto the spatial gene expression datasets confirmed the colocalization of monocyte/macrophages with the IL-1β response gene signature (Fig. 5e). Similar analyses of an independent RA sample confirmed these results (Extended Data Fig. 5a-e). This association was further validated by an unbiased correlation of the FLS states, cytokine response gene signatures, and cell type specific gene signatures within individual spots that showed a correlation between monocytes/macrophages, lining FLS and the IL-1β response gene signature (Extended Data Fig. 5f). These results suggest that cytokine signaling shapes multiple spatially distinct microenvironments within the inflamed RA synovium with IL-1β from either resident macrophages or infiltrating monocytes defining the synovial lining FLS.

**Figure 5.**
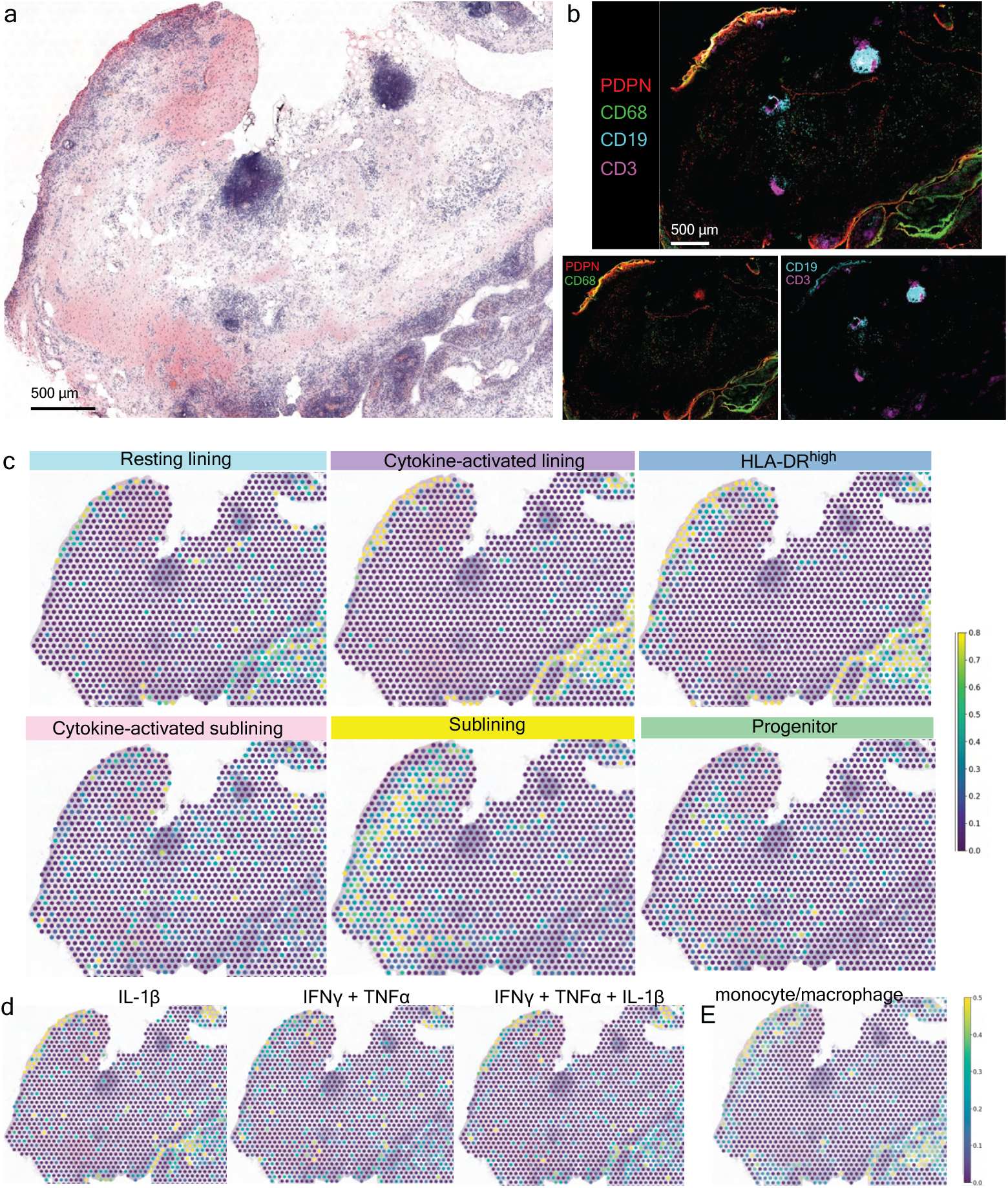
Cytokine signaling is spatially constrained and correlated with cellular localization. **a,** H&E staining of a representative tissue section used for ST (patient 4 in Table S1). **b,** IF image from a serial tissue section directly adjacent to that used for ST analysis. The down staining is for the following markers: PDPN (red), CD68 (green), CD19 (cyan), and CD3 (magenta). (N = 2 tissues for ST with 2 adjacent sections each) **c,** Relative expression of FLS states defined in Fig. 1g in each RNA capture area on the ST slide. **d,** Relative expression of FLS cytokine response signatures in each RNA capture area on the ST slide. **e,** Relative expression of synovial macrophage specific gene signature from Zhang et al^2^ in each RNA capture area on the ST slide.

## Discussion

Phenotypic and functional heterogeneity of parenchymal cells within a given tissue depends on both constant cell-intrinsic differentiation programs and cell extrinsic cues afforded by stable and transient interactions with tissue resident and infiltrating cells. The latter are represented for the most part by diverse types of immune cells, which, when activated, produce cytokines and other mediators that can act on the parenchymal cells changing the range of their physiological or homeostatic states. In RA, the synovium, which is in health a non-barrier immunologically quiescent tissue, experiences a massive influx of both innate and adaptive immune cells. In this setting, FLS experience differentiation signals, such as the endothelium-derived Notch signaling gradient that shapes FLS in the synovial sublining region^8^, and coincident exposures to multiple cytokine and other inflammatory mediators. Using paired scRNA/ATAC-seq and ST analyses assisted by *in vitro* generated cytokine response signatures, we demonstrated that leukocyte-derived cytokines play a key role in the formation of discrete, spatially defined, but likely dynamic FLS states with distinct inferred functionality. We also observed, contrary to a reported potentiation of inflammatory cytokine responses in macrophages^22^, Notch activation in FLS attenuated cytokine responses. This finding may explain the relatively dampened cytokine response gene signatures observed in the sublining FLS state, which was distinguished by the highest *NOTCH3* expression, as compared to the cytokine-activated sublining FLS state.

Our analysis of rich paired datasets of single cell transcriptomes and epigenomes allowed us to detect previously unappreciated heterogeneity of FLS in highly inflamed RA synovium, including an inflamed sublining state driven by IFNγ and TNFα and two lining states defined by differential cytokine responses. The observation that the cytokine-activated lining FLS transcriptome was enriched for genes downstream of IFNγ, TNFα and IL-1β was consistent with the expression of the latter two cytokines by synovial macrophages^2^, and the coincident positioning of macrophages with the local IL-1β signature in the synovial lining in our ST analysis. Potential sources of IFNγ and STAT1 activation within the synovial lining remained less clear. While CD8^+^ T cells have been described as the dominant producers of IFNγ within the synovium, it is possible that other cells such as NK cells^3^ or myeloid cells^25^ may contribute. Finally, it is probable that the prominent pSTAT1 signal observed in the synovial lining FLS (Fig. 4c), which has also been reported previously^26^, is the result of the action of alternative drivers of STAT1 activation including type I IFN^26^, whose expression can be driven by IL-1β^27^.

The prominent IL-1β response signature in the cytokine-activated lining FLS state is notable given its possible functional and therapeutic implications. First, IL-1β is the primary inducer of MMPs, which have been implicated in FLS invasiveness^28^. This invites the possibility of a functional dichotomy between sublining and lining FLS in RA parallel to that observed in an experimental arthritis model in mice, where the lining FLS are uniquely responsible for destruction of cartilage and bone^29^. Blocking IL-1 may antagonize the capacity of FLS to assume this MMP expressing lining state and, thus it is possible that for a subset of patients the addition of IL-1 inhibition during flares could prove effective for preventing joint destruction. Second, the cytokine-activated lining FLS state may drive migration of neutrophils into the synovial fluid, where there is a surfeit of neutrophils during RA flares. In this regard, the inflammatory cancer-associated fibroblasts in colorectal adenocarcinomas, which we found to share extensive transcriptional similarity with the cytokine-activated lining FLS state, express neutrophil chemoattractants including *CXCL1* and *CXCL8* and their location was spatially correlated with the accumulation of neutrophils^11^. In RA, IL-1β produced by macrophages in the lining region likely drives the high expression of *CXCL1* observed in the cytokine-activated lining FLS. Finally, the prominence of the cytokine-activated lining FLS state observed in highly inflamed RA synovium may have prognostic implications. A recent study of pathotypes in inflammatory bowel disease showed an association of an IL-1β-activated fibroblast signature with a lack of response to multiple therapies^30^.

Besides the cytokine-driven FLS states, we defined a CD34^high^THY1^+^PDPN^+^ progenitor-like FLS population that was devoid of cytokine response signatures and shared extensive transcriptional similarity with fibroblast populations found in both the colon and skin. The abundance of this progenitor-like population appears to vary between synovial tissues of individual patients and may depend on disease activity or treatment. Future studies will help determine if the paucity of cytokine response gene signatures in these FLS owes to their sequestration from cytokine-producing immune cells, or cell-intrinsic attenuation of cytokine signaling.

In conclusion, we established a spatial atlas of heterogeneity of synovial fibroblast states in RA defined by their distinct transcriptional signatures and patterns of chromatin accessibility driven by differential local exposure to immune cell-derived pro-inflammatory cytokines. The resulting datasets will serve as a rich resource for future investigation of RA pathogenesis through integration of unique and shared characteristics of inflamed synovial fibroblasts described herein with other disease-associated signals, such as complement activation^31^, and potential antigen presentation via HLA-DR.

## Acknowledgments

We thank HSS orthopedic surgeons, clinical research coordinators (particularly Diyu Fisher and Edoardo Spolaore) and the HSS patients who contributed to this study. We acknowledge the Accelerating Medicines Partnership® (AMP®) in Rheumatoid Arthritis and Lupus Network for the stimulating discussions and the large-scale sequencing of arthritis patient synovial tissues that formed the basis for this study. AMP is a public-private partnership (AbbVie Inc., Arthritis Foundation, Bristol-Myers Squibb Company, Foundation for the National Institutes of Health, GlaxoSmithKline, Janssen Research and Development, LLC, Lupus Foundation of America, Lupus Research Alliance, Merck Sharp & Dohme Corp., National Institute of Allergy and Infectious Diseases, National Institute of Arthritis and Musculoskeletal and Skin Diseases, Pfizer Inc., Rheumatology Research Foundation, Sanofi and Takeda Pharmaceuticals International, Inc.) created to develop new ways of identifying and validating promising biological targets for diagnostics and drug development.

## Author contributions

MHS conceived the project, collected clinical data, designed experiments, performed experiments, prepared figures, wrote the manuscript.

VRG analyzed sequencing data, prepared figures and edited the manuscript.

MS analyzed sequencing data.

AK performed experiments.

EFD scored histology slides.

SMG collected clinical data.

TMN designed experiments, supervised research experiments, analyzed sequencing data and prepared figures.

CSL oversaw the computational analysis and edited the manuscript.

LTD conceived the project, acquired funding, designed experiments, supervised research experiments, oversaw data analysis and edited the manuscript.

AYR conceived the project, acquired funding, designed experiments, supervised research experiments, oversaw data analysis and wrote the manuscript.

## Code availability

The customized code used in the present study is publicly available at: https://github.com/viannegao/RA_Fibroblast_Analysis

## Data availability

All sequencing data generated in this paper were deposited in the Gene Expression Omnibus (GEO) under accession numbers: (accession number to be generated)

All materials and data transferred between HSS and MSKCC were covered under material transfer agreements.

---

**Extended Data Figure 1:**
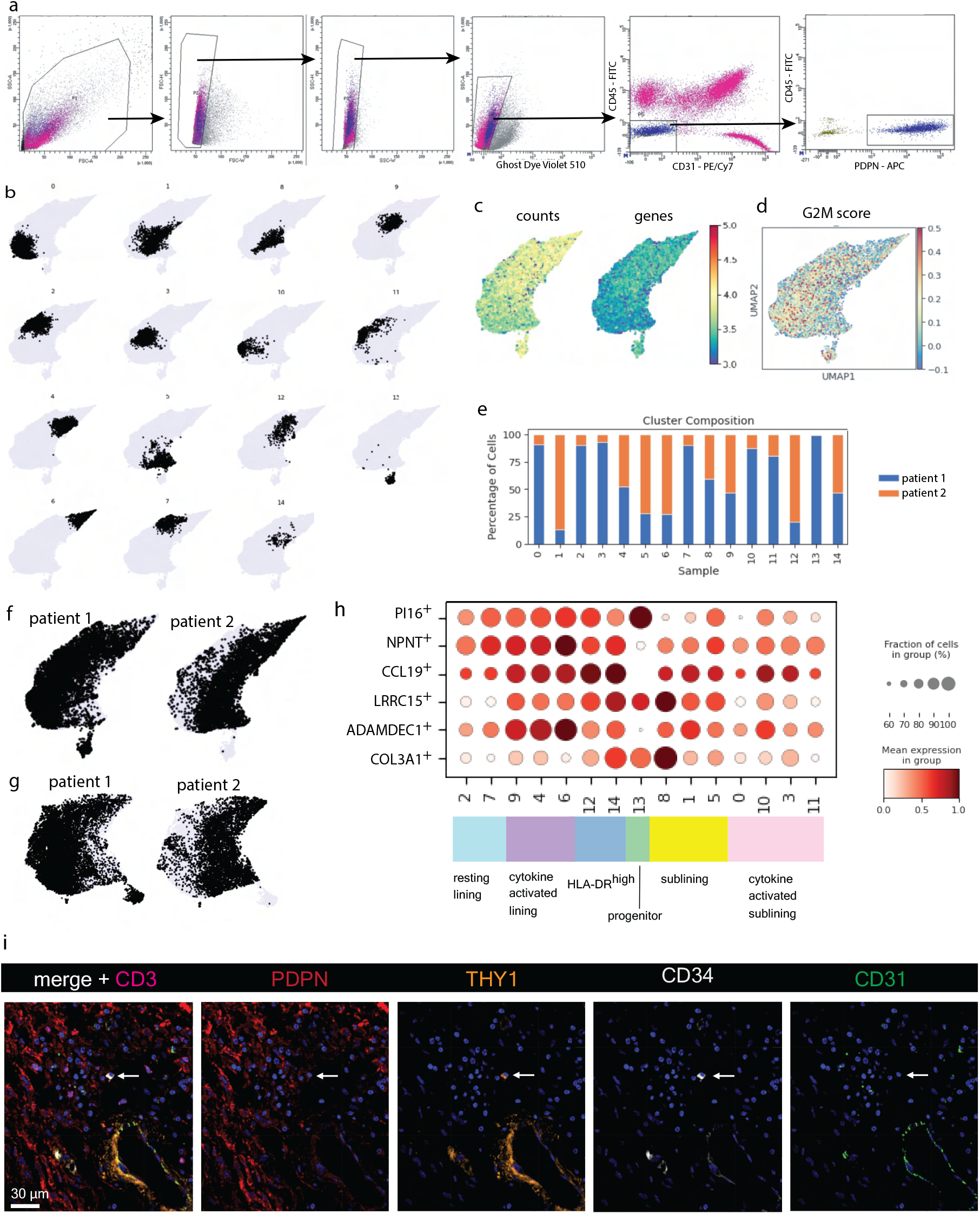
scRNA-seq analysis of FLS isolated from inflamed RA synovium. **a,** Gating strategy for FACS sorting of FLS. **b,** Individual FLS clusters shown on UMAP. **c,** Numbers of UMI counts and unique genes expressed in each cell shown on UMAP. **d,** Cell cycle G2M score per cell shown on UMAP. **e,** Cluster composition by patient. **f,** Distribution of cells from each patient without batch correction. **g,** Distribution of cells from each patient with MNN batch correction. **h,** Dot plot with relative expression of fibroblast signatures from all clusters defined by the human perturbed fibroblast atlas from Buechler et al^9^. **i,** Representative confocal image of PDPN (red), THY1 (orange), CD34 (white), CD31 (green), CD3 (magenta) and nuclear marker (blue) from RA synovial tissue (N = 4 tissues). White arrow indicates individual CD34^+^, THY1^+^, PDPN^+^ cell not within the vasculature.

**Extended Data Figure 2:**
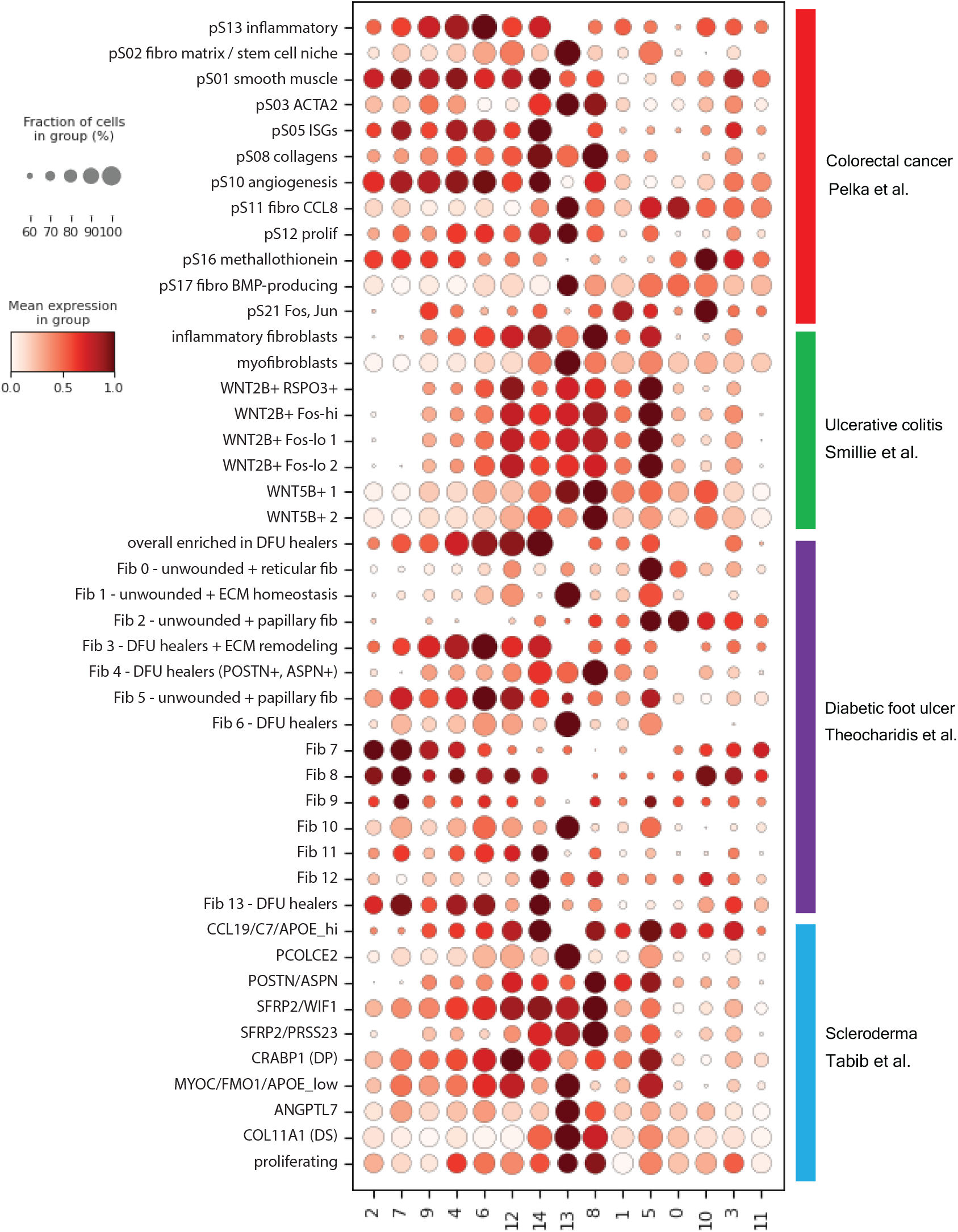
Expression of non-synovial fibroblast gene signatures in FLS. Dot plot showing relative expression of all gene signatures from published tissue fibroblast populations^11–14^ in FLS clusters shown in Fig. 1a colored according to FLS states defined in Fig. 1g.

**Extended Data Figure 3:**
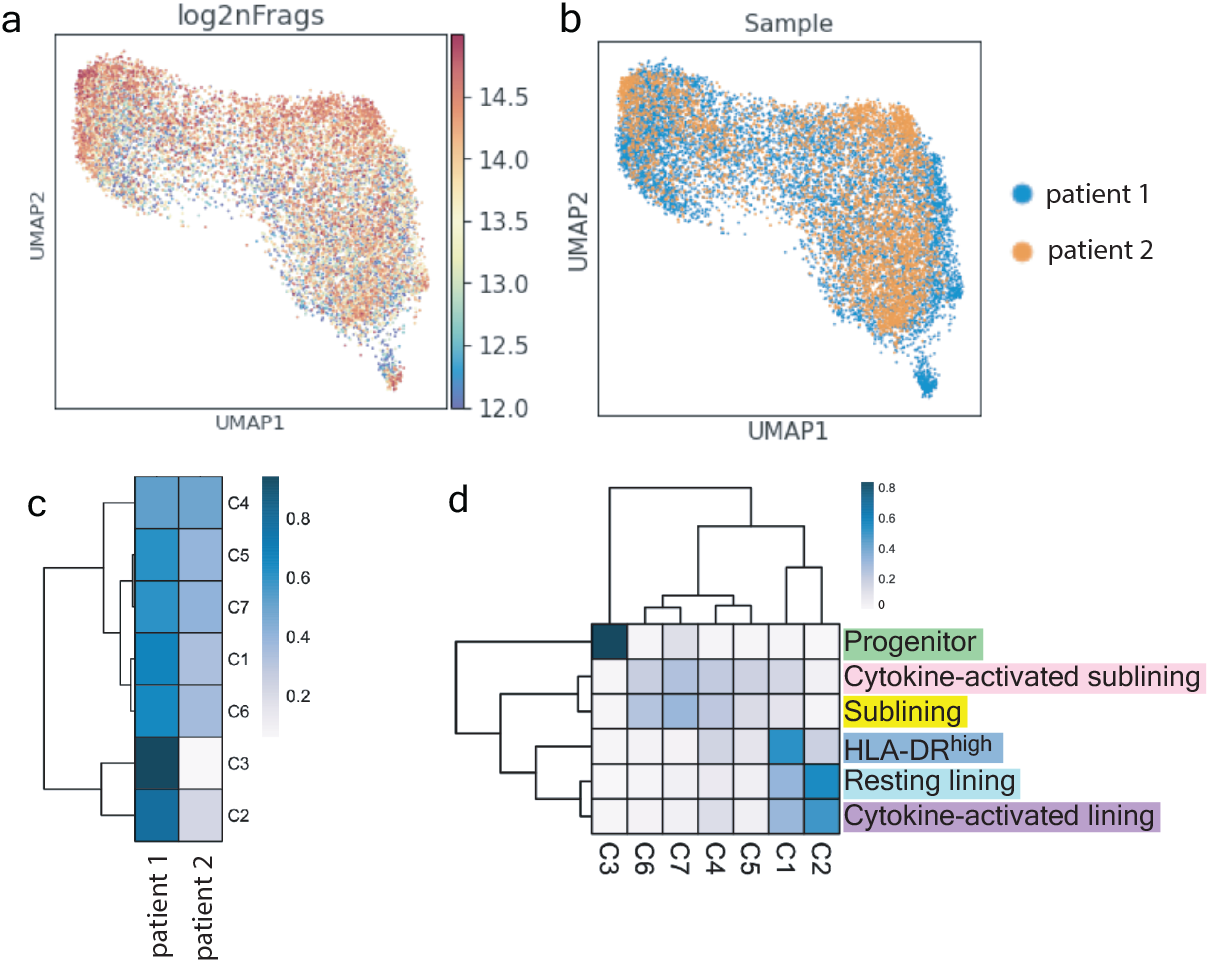
Paired scATAC-seq analysis of isolated FLS. **a,** Logged number of fragments detected in each cell shown on UMAP. **b,** Distribution of cells from each patient. **c,** scATAC-seq cluster composition by patient. **d,** scATAC-seq cluster composition by FLS states.

**Extended Data Figure 4:**
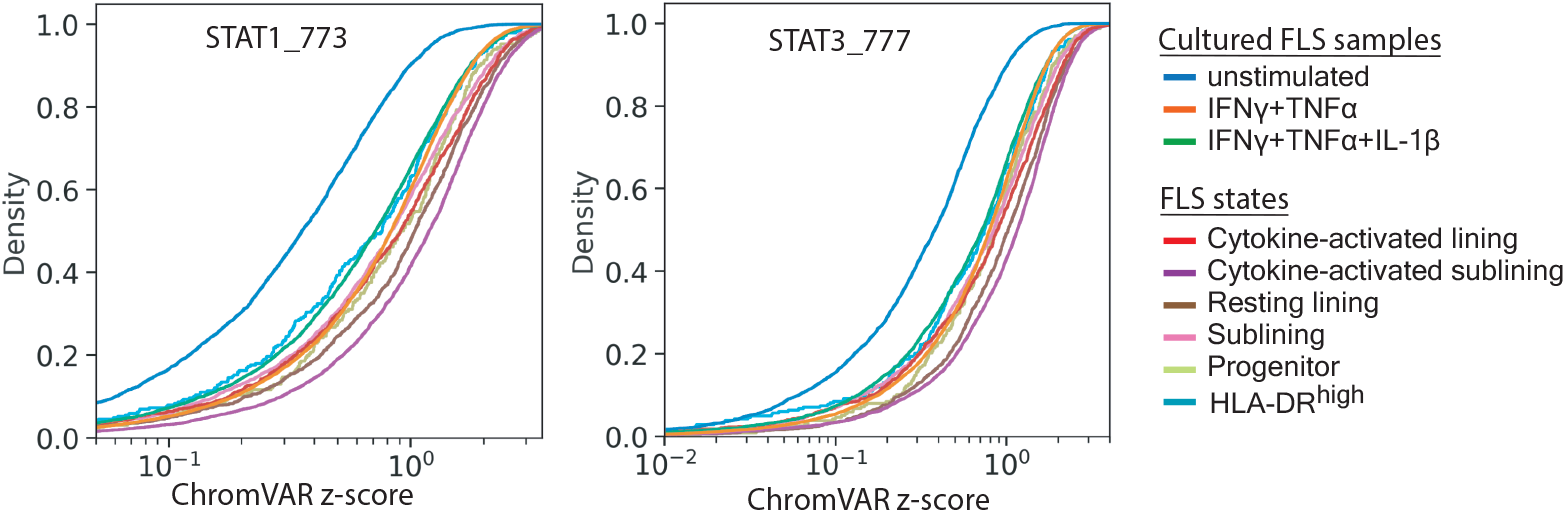
Dynamics of chromatin accessibility of STAT motif containing cis-regulatory elements across *in vivo* FLS states and *in vitro* cytokine stimulated or unstimulated FLS. Empirical cumulative distribution function (ECDF) 600 plots of ChromVAR z-scores for STAT motifs from scATAC-seq analysis of the indicated FLS samples or states are shown.

**Extended Data Figure 5:**
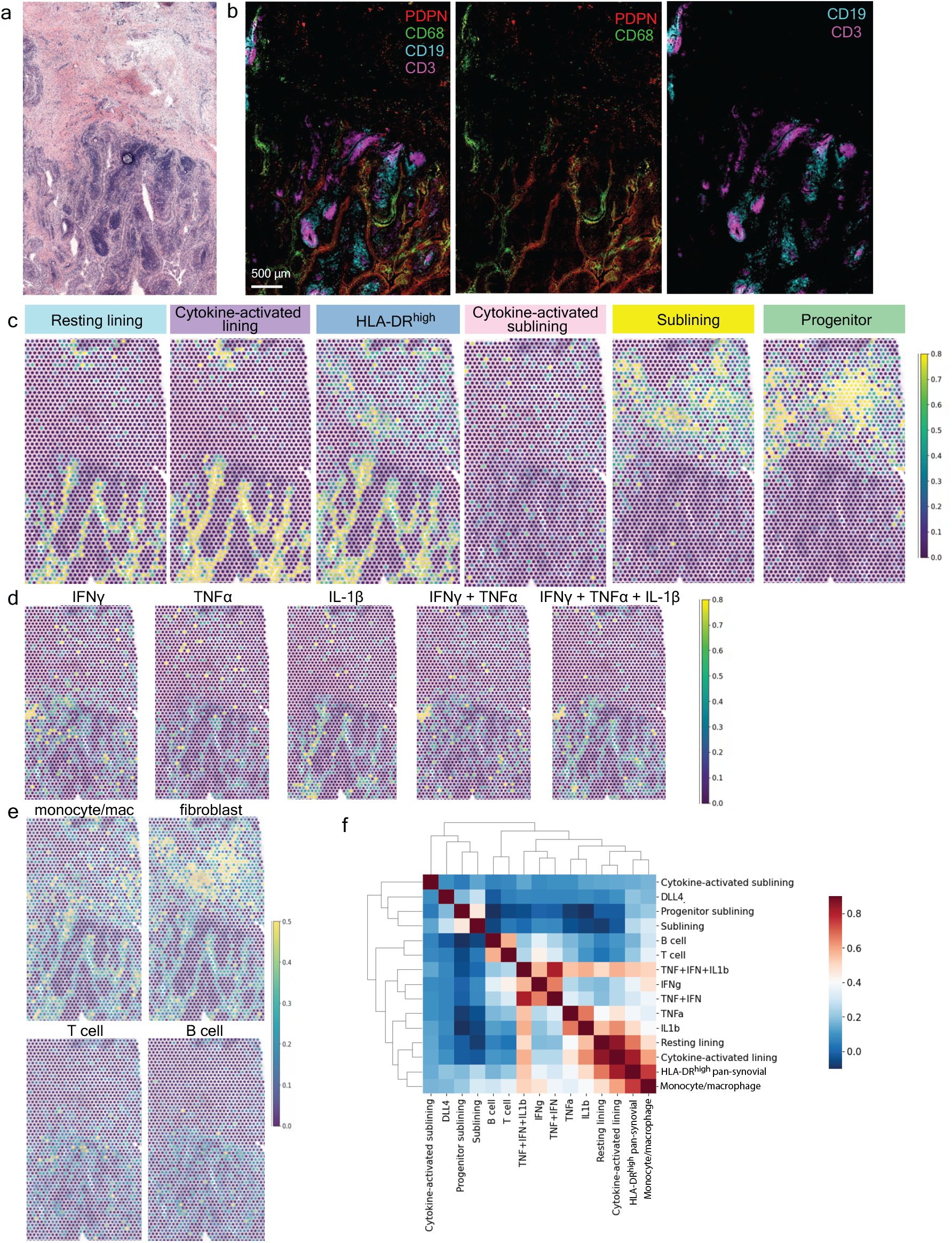
Spatial transcriptomic analysis of inflamed RA synovial tissue. **a,** H&E staining of a tissue section used for ST (patient 3 in Table S1). **b,** IF image from a serial tissue section directly adjacent to that used for ST stained for: PDPN (red), CD68 (green), CD19 (cyan), and CD3 (magenta). **c,** Relative expression of gene signatures of FLS states defined in Fig. 1g in each RNA capture area on the ST slide. **d,** Relative expression of FLS cytokine response gene signatures in each RNA capture area on the ST slide. **e,** Relative expression of synovial cell type specific gene signatures from Zhang et al^2^ in each RNA capture area on the ST slide. **f**, Correlation between FLS state gene signatures derived from scRNA-seq data (from c), *in vitro* cytokine response gene signatures (from d), and cell type specific gene signatures (from e) within individual RNA capture spots.

## Supplementary Information

**Table S1**: Patient characteristics for synovial tissue samples used in this study. Samples from patients 1 and 2 were used for the paired scRNA and scATAC sequencing. Samples from patients 3 and 4 were used for spatial transcriptomics.

**Table S2**: Differentially expressed genes for clusters defined in Fig. 1a and states defined in Fig. 1g.

**Table S3**: GSEA results for each state defined in Fig. 1g.

**Table S4**: Differentially expressed genes for RA FLS stimulated with cytokines in vitro.

**Table S5**: Genes differentially expressed with the addition of Notch ligand DLL4 to cytokine stimulation.

## Methods

### Human synovial tissue

Synovial tissue was obtained from patients consented into the HSS FLARE study of RA patients undergoing arthroplasty or synovectomy (IRB no 2014-233). On day of surgery, samples were cryopreserved in as small fragments in CryoStor CS10 (Stem Cell Technologies #07959). Synovial tissue quality and grading of synovitis^32^ were evaluated by histologic analysis (H&E).

### Sample preparation for single cell sequencing

Synovial tissue samples were disaggregated into a single-cell suspension as described previously^2^. Briefly, fragments were minced and enzymatically digested (Liberase TL (Sigma-Aldrich) 100 μg/mL and DNaseI (New England Biolabs) 100 μg/mL in RPMI) for 30 min at 37°C. Disaggregated cells were assessed for quality and viability (Nexcelom Cellometer Auto 2000) and then stained with antibodies to CD45 (2D1), CD31 (WM59), PDPN (NZ-1.3) and Ghost Dye Violet 510 (Tonbo) for fluorescence activated cell sorting (BD FACSAria III Cell Sorter). Synovial fibroblasts (CD45^-^, CD31^-^, PDPN^+^ were collected directly into FACS buffer. Individual nuclei were prepared using the 10x Genomics protocol CG000365-Rev A. Nuclei were submitted for sequencing via Chromium Single Cell Multiome ATAC + Gene Expression (10x Genomics) by the Integrated Genomics Operation core facility at the Sloan Kettering Institute.

For cultured cytokine stimulated FLS, synovial tissues were dissociated into single cells as above, cultured in MEM alpha (ThermoFisher Scientific Gibco 12561056) with 10% Fetal bovine serum (R&D systems S11550) as well as 1% penicillin/streptomycin (ThermoFisher Scientific 15070063) and 1% L-glutamine (ThermoFisher Scientific 25030081). Cells were passaged using TrypLE Express Enzyme (ThermoFisher Scientific Gibco 12605010) until a FLS monoculture was present (>3 passages). At passage 4 with TNFα (20 ng/mL) + IFNγ (5 ng/mL) or TNFα (20 ng/mL) + IFNγ (5 ng/mL) + IL-1β (1 ng/mL) for 24 hours prior to harvesting and isolating nuclei as above.

Cytokine sources: recombinant human TNFα from PeproTech (#300-01A), recombinant human IFNγ from Roche (#11040596001), recombinant human IL-1β from PeproTech (#200-01B).

### Quantification and Statistical Analysis

#### Pre-processing of single cell multiome ATAC + gene expression data

RNA and ATAC libraries for each patient were aligned using cellranger-arc software (v1.0.0, 10x Genomics) against 10x genomics reference refdata-cellranger-arc-GRCh38-2020-A using default parameters. The output files fragments.tsv.gz and filtered_feature_matrix.h5 were utilized for downstream processing and quality control analysis. We then perform the following additional cell filtering steps: 1) cells with a high fraction of mitochondrial molecules were filtered (> 20%); 2) clusters resembling contaminating immune cell populations were removed and 3) clusters with low library complexity were filtered (cells that express very few unique genes). Putative doublets were removed using the DoubletDetection package (https://doi.org/10.5281/zenodo.2658729). Cells or nuclei that passed these QC cutoffs were used to generate sparse count matrices and filtered fragments.tsv.gz files for downstream analysis.

### Single-cell RNA-seq data analysis

#### Preprocessing, dimensionality reduction, clustering

Combining the two patient samples yielded a filtered count matrix of 15736 cells by 36391 genes, with a median of 4156 molecules per cell. The count matrix was then normalized by library size and scaled to 100,000 per cell for analysis of the combined dataset. Highly-variable genes were identified using the Scanpy highly_variable_genes function with batch_key=‘sample’. Principal component analysis (PCA) was performed on the normalized expression of highly-variable genes with the top 30 principal components (PCs) retained. We first performed clustering on the combined dataset using Phenograph with k = 30 to identify 15 clusters. To aid subtype annotation, we merged these clusters into meta-clusters based on the correlation in cluster mean expression of highly-variable genes. We then annotated these meta-clusters based on enriched gene pathways identified by GSEA and previously-published datasets (Alivernini et al^4^ and Zhang et al^2^). To evaluate the amount of batch effect between the two patients, we performed mutual nearest neighbor correction using Scanpy’s mnn_correct function. Specifically, we limited the analysis to only the highly-variable genes and used svd_dim = 50.

#### Visualization of single-cell RNA-seq

To visualize single cells of the two patients, we used UMAP projections(McInnes et al., 2018) to generate lower dimensional representations using knn = 30 and min_dist = 0.2.

#### Differential expression in scRNA-seq

We performed differential expression for the following comparisons: 1) samples from each cytokine-stimulation conditions vs samples from non-stimulated, cultured samples (Tables S4), 2) each fibroblast state vs rest (Table S2 tab 2), and 3) each unsupervised cluster vs rest (Table S2 tab 1). All differential expression was performed using MAST (version 1.8.2)^33^, which provides a flexible framework for fitting a hierarchical generalized linear model to the expression data. We used a regression model that adjusts for cellular detection rate (cngeneson, or number of genes detected per sample):

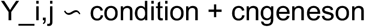

where condition represents the condition of interest and Y_i is the expression level of gene i in cells in cluster j, transformed by natural logarithm with a pseudocount of 1.

To homogenize cell sampling per condition, we downsampled such that the cell complexity (i.e. the number of genes per cell) was evenly matched across groups. We partitioned cells from each group into 10 equally sized bins based on cell complexity and subsampled from each bin to match cell complexity distribution across samples. We downsampled to at most m cells per group, where m is the median number of cells per group. We considered genes to be significantly differentially expressed for Bonferroni-adjusted p-value < 0.05.

#### Identifying enriched gene pathways in single-cell RNA-seq data

Enriched gene pathways were identified using pre-ranked GSEA, as implemented by the R package fGSEA^34^ using 10,000 permutations. Gene ranks were calculated using −log(p-value)*log fold change based on MAST^33^ differential expression. To assess enriched pathways in clusters, we used HALLMARK and KEGG subset of Canonical Pathways in MSigDB v 7.1^35^. We considered pathways with Benjamini-Hochberg adjusted p-values < 0.1 to be significant.

#### Scoring Gene Signature Expressions

To score the single-cell expression of gene signatures, we first transformed the library size-normalized, log-transformed data by z-score and calculated the average expression of each curated gene set per cell type subtracted from the average expression of a reference set of genes using the score_genes function in scanpy. The subsequent cell type scores were transformed again by z-score. For comparisons to published datasets, we used the top 30 genes after sorting by adjusted p value (padj) for top differentially expressed genes for each unsupervised cluster/cell type.

#### Correlating Spatial Gene Signature Expression

To correlate the spatial localization of gene signatures of interest, we computed the pearson correlation coefficient of their expressions across individual spots for all samples combined. The signatures are generated as described above.

### Single-cell ATAC-seq data analysis

#### Preprocessing, dimensionality reduction, clustering

We preprocessed the filtered fragments.tsv file using the ArchR package^36^ v1.0.1. Specifically, we binarized sparse accessibility matrices binned at 500bp tiles across the genome. We then perform iterative latent semantic indexing (LSI) on the tile matrix to generate 30 components. For downstream analysis, we filter 3 components strongly correlated (cor > 0.5) with the number of fragments detected per cell, as well as one component that is strongly correlated with batch. For visualization, we used the addUMAP function in ArchR with the following parameters: nNeighbors = 150, minDist = 0.05, metric = cosine. Clustering was done using the addClusters function in ArchR with the following parameters: method = “Seurat”, knnAssign = 50.

#### Peak-calling and TF motif accessibility scoring

Filtered fragments for cells in each sample were aggregated and used as input to the MACS2 peak caller^37^; parameters -f BED, -g 2.7e9, --no-model, --shift -75, --extsize 150, -q 0.05). Peaks are filtered using an IDR cutoff of 0.05. We subsequently added motif annotations using “addMotifAnnotations” with the CisBP motif database and computed chromVAR deviations for each single cell with “addDeviationsMatrix”.

#### Identifying enriched motifs per cluster

To identify differentially accessible motifs for each group of interest, we used the rank_genes_groups function in scanpy with method=‘wilcoxon’ and corr_method = ‘benjamini-hochberg’ on the chromVAR zscore matrix. Motifs were filtered to include only those for which the corresponding transcription factor was expressed by >20% of cells in the corresponding FLS state. The top 10 motifs after ranking by ‘score’ were then selected for plotting in the heatmap in Fig. 3d.

### In vitro FLS culture and stimulation for bulk RNA sequencing

Synovial tissues from 4 donors (RF^+^ and/or CCP^+^) were disaggregated and cultured as above. At passage 4 or 5, cells from the 4 donors were pooled and were plated into 12 well plates at 70,000 cells/well. Cells were allowed to adhere and were then stimulated with TNFα (0.1 ng/mL), IFNγ (0.05 ng/mL), or IL-1β (0.01 ng/mL) alone or in combination for 24 hours in triplicate. For DLL4 treatment, cell culture plates were coated with recombinant DLL4-FC (R&D systems) overnight at 4°C at 0.5 μg/mL prior to the addition of FLS and cytokines. Concentrations for all stimuli were determined based on an initial titration experiment with 4 concentrations per stimulus (10x dilutions starting with 100 ng/mL TNFα, 50 ng/mL IFNγ, 10 ng/mL IL-1β and 5 μg/mL DLL4) and bulk 3’ RNA sequencing to determine the number of differentially expressed genes relative to untreated cells. After stimulation, we lysed cells, isolated RNA (Zymo Research R1052), prepared libraries (Lexogen QuantSeq 3’ mRNA-Seq Library Prep Kit (FWD) for Illumina 015.96) and the Integrated Genomics Operation at the Sloan Kettering Institute sequenced samples (bulk RNA sequencing).

### Stimulated FLS bulk RNA sequencing data analysis

Reads from 3’ RNA sequencing of fibroblasts treated with cytokines were processed using version 2.5.3 of the snakePipes mRNA-seq pipeline^38^ using the flags “--reads ‘_R1_001’ ‘_R2_001’ --mode ‘alignment’ --trim --trimmerOptions ‘-a A{10}N{90}’“. In brief, this pipeline trims reads using Cutadapt, aligns them using STAR to the genome (release 34 of GRCh38 with Gencode annotations), and then aggregates gene-level counts using featureCounts. Differentially expressed genes for each condition were then defined relative to control cells using DESeq2. Genes that were up- or downregulated at *p* < 0.05 following correction for multiple hypothesis testing for each single cytokine treatment were used to define expression signatures for each cytokine. The distributions shown in Fig. 4b are of the (shrunken) log2 fold change estimates of these genes relative to control cells estimated by DESeq2 in cells treated with the indicated cytokines or combinations of cytokines.

### Multicolor immunofluorescence

Synovial tissue was fixed in 1:4 dilution Fixation/Permeabilization solution (BD Biosciences Cytofix/Cytoperm Cat No. 554714) in PBS pH 7.4 for 16-20 hours at 4°C. Tissue was washed 3x with PBS then placed in 30% sucrose in 0.1M sodium phosphate buffer pH 7.4 until the tissue fell to the bottom of the tube (~6 hrs) at which point tissue was embedded in optimal cutting temperature compound (OCT), frozen on dry ice and stored at −80°C until sectioning (10 μm thick). For staining, tissues were rehydrated on slides, permeabilized with 0.1% triton in PBS for 10 min and blocked with 5% normal goat serum (ThermoFisher Scientific 31873) in PBS for 45 min prior to staining with primary antibodies (5 hrs RT or 21 hrs at 4°C) followed by secondary antibodies (2 hours RT). Appropriate isotype controls were used on a separate section. After antibody stains, slides were washed, stained for nuclei (DAPI – ThermoFisher Scientific 62248 – 1:2000 for 5 min RT) and mounted with Fluoromount G (ThermoFisher 00-4958-02). Images were acquired with a Leica SP8 confocal microscope (40x oil immersion). Image analysis (merging of channels) was performed with Imaris cell imaging software.

### Spatial Transcriptomics

Fresh synovium was immediately embedded in OCT and frozen using isopentane cooled by liquid nitrogen. We used Visium Spatial Gene Expression platform (10x Genomics) in conjunction with the Integrated Genomics Operation and Molecular Cytology core facilities at the Sloan Kettering Institute. For this, tissue was sectioned (10 μm sections, 2 tissue sections in 2 replicates each per slide, capture area 6.5 x 6.5 mm), stained with H&E, and permeabilized. This was followed by cDNA library construction with spatial barcoding and sequencing.

### Spatial transcriptomics data analysis

#### Preprocessing, dimensionality reduction

Spatial sequencing data from 2 patients (2 samples each) were aligned using the Space Ranger (v1.2.2, 10x genomics) pipeline to the 10x genomics reference genome refdata-gex-GRCh38-2020-A using default parameters to derive a feature spot-barcode gene expression matrix. Combining the 4 samples yielded a filtered count matrix of 12257 spots by 19809 genes, with a median of 1754 molecules per spot. The count matrix was then normalized by library size and scaled to the median of total counts of all cells before normalization for analysis of the combined dataset. We then natural-log transformed the reads with a pseudocount of 1. Seurat-v3.2 package was then used to select top variable genes for spatial RNA-seq clustering. Highly-variable genes were identified using the Scanpy highly_variable_genes function with batch_key=‘sample’ and n_top_genes = 2000. PCA was performed on the normalized expression of highly-variable genes with the top 50 principal components (PCs) retained.

#### Scoring Gene Signature Expressions

To score the single-cell expression of gene signatures, we further transformed the data by z-score and calculated the average expression of each curated gene set per cell type subtracted from the average expression of a reference set of genes using the score_genes function in scanpy. The subsequent cell type scores were transformed again by z-score. Gene signature expressions were visualized using the scanpy.pl.spatial function.

### Antibodies used

Antibody (vendor; catalog number; clone; lot; dilution – final concentration)

#### For sorting FLS

Anti-CD45-FITC (eBiosciences; 11-9459-42, 2D1; lot 4271593; 1:100)
Anti-PDPN-APC (Invitrogen; 17-9381-42; NZ-1.3; lot 1988690; 1:100)
Anti-CD31-PE/Cy7 (Biolegend; 303118; WM59; lot B276836; 1:100)
Ghost Dye Violet 510 (Tonbo; 13-0870-T100; no clone; lot D0870040521133; 1:1000)

#### For immunofluorescence

Primary:

Anti-PDPN (Invitrogen; 14-9381-82; NZ-1.3; lot 2400405; 1:100 – final 5 μg/mL)
Anti-HLA.DR-AF488 (Biolegend; 307620; L243; lot B271228; 1:100 – 2 μg/mL)
Anti-CD3-BV480 (BD biosciences; 566105; UCHT1; lot 0079903; 1:100)
Anti-CD8-AF647 (Biolegend; 344725; SK1; lot B270006; 1:50 – final 1 μg/mL)
Anti-pSTAT1-PE (Biolegend; 686403; A15158B; lot B327686; 1:50 – final 0.12 μg/mL)
Anti-CD68-AF488 (Biolegend; 333812; Y1/82A; lot B278908; 1:10 – final 2.4 μg/mL)
Anti-CD19-PE (Biolegend; 302208; HIB19; lot B273506; 1:20 – final 2.5 μg/mL)
Anti-CD90-AF700 (R&D systems; FAB2067N; Thy1A1; lot 1569061; 1:50 – final 4 μg/mL)
Anti-CD34-AF647 (Biolegend; 343507; 581; lot B312791; 1:100 – final 2 μg/mL)

Secondary:

Anti-rat-AF594 (Biolegend; 405422; polyclonal; lot B302011; 1:1000)

