## Supplemental Table 1 for "Heterogeneity of Inflammation-associated Synovial Fibroblasts in Rheumatoid Arthritis and Its Drivers"

Table S1

| | Patient 1 | Patient 2 | Patient 3 | Patient 4 |
|---|---|---|---|---|
| RA criteria met | 1987 and 2010 | 1987 and 2010 | 1987 and 2010 | 2010 |
| Disease duration (yrs) | 11 | 10 | 23 | 4 |
| CCP | > 3x ULN | > 3x ULN | > 3x ULN | > 3x ULN |
| RF | 2x ULN | > 3x ULN | > 3x ULN | negative |
| Active treatment | Methotrexate | NSAIDS | Leflunomide, adalimumab | Sulfasalazine, hydroxychloroquine |
| Prior treatments | TNF inhibitor | Methotrexate, TNF inhibitor | Methotrexate | Methotrexate, abatacept, adalimumab, certolizumab, infliximab, tocilizumab, tofacitinib |
| Joint | Knee | Knee | Wrist | Knee |
| Krenn Score | 8 | 5 | 6 | 4 |
| Pathotype | lymphoid | lymphoid | lymphoid | lymphoid |
| Lining thickness | 4-5 cells | > 5 cells | > 5 cells | 4-5 cells |
| Stromal cellularity | Greatly increased | Normal | Normal | Normal |
| Inflammatory infiltrate | Dense band-like or numerous large follicle-like aggregates | Numerous lymphocytes or plasma cells, sometimes forming follicle-like aggregates | Dense band-like or numerous large follicle-like aggregates | Numerous lymphocytes or plasma cells, sometimes forming follicle-like aggregates |
